# Hypothalamic Interleukin 6 linked to sex-specific behavioral deficits following adolescent social isolation

**DOI:** 10.64898/2026.04.16.719013

**Authors:** Chetan Mishra, Arundhati Gupta, Beena Pillai, Arpita Konar

## Abstract

Social isolation refers to an extreme form of social deprivation that has enduring effects on the brain and behavior. Adolescents show selective vulnerability to such heightened social stress, displaying aberrant behavior and psychiatric ailments. The post-weaning social isolation rodent model has been widely used to recapitulate such behavioral anomalies and delineate their mechanistic bases. Here, we aim to identify how prolonged social isolation during adolescence affects neuroimmune responses in both sexes and the implications for behavioral outcomes, particularly aggression. While males subjected to adolescent isolation were hyper-aggressive with pathological signs, females showed reduced social exploration and inactivity. Cytokine profiling in core brain regions implicated in aggression revealed reduced interleukin 6 (IL6) levels, specifically in the hypothalamus, in both sexes. Other proinflammatory cytokines, including interferon-gamma and interleukin-1beta, were unaltered. IL6-responsive genes, SOCS3 and TIMP1, were also downregulated in the hypothalamus of both socially isolated males and females. The hypothalamus is crucial for stress responsiveness and the expression of excessive aggression. Despite behavioral dimorphism, reduced IL6 levels in both sexes may indicate differences in downstream signaling and roles beyond classical immune responses. Our findings suggest that hypothalamic IL6 may be a key mediator of adolescent social isolation, which is associated with aberrant behavior, including aggression.

## Introduction

Social behavior is fundamental to survival, reproductive fitness and development of the animal kingdom. Chronic physical ailments, environmental barriers, social stigma, and age transitions are key factors imposing social isolation in humans and a huge mental health burden. Social isolation emerged as a massive psychosocial stressor when Covid-19 hit the globe and precipitated long-term psychiatric illnesses such as mood disorders, cognitive deficits, aggression and impulsivity^1,2^. Previous studies have shown that social stress during adolescence is a prominent risk factor for a multitude of psychopathologies in later life. Anxiety^3,4^, depression^5,6^, bipolar disorder^7^, schizophrenia^8^, aggression^9,10^, suicidal ideation^6^, and self-harm^11^ have been linked with adolescent stress, though the biological basis remains poorly understood.

Adolescence is characterized by major brain-body transitions, including increased peer influence, a sense of independence, and the exploration of sexual preferences^12^. The brain undergoes dynamic reorganizations during this crucial period, including grey and white matter integrity^13,14^, maturation of the PFC^15,16^, and structural and functional changes in GABAergic and glutamatergic neurotransmission^17^. PFC-mediated decision-making and impulsivity undergo synaptic pruning and dendritic spine alterations during adolescence^18^, which are essential for adapting to the social environment. However, stressors like isolation can disrupt such neural remodeling increasing vulnerability to behavioral aberrations and mental health challenges. The hypothalamus, a key regulatory region of the endocrine system, also plays a role in the transition from childhood to adolescence^19,20^. Because the hypothalamic-pituitary axis is central to stress responsiveness and cortisol release, early-life social stress can impair hypothalamic-regulated motivated behaviors, such as aggression, mating, and feeding.

Our immune system parallels radical shifts during adolescence^21,22^ along with rewiring of neural circuits. Accumulating studies highlight the role of neuroimmune interactions in brain plasticity, social stress sensitivity and behavioral outcomes^23,24^. Chronic stress models have shown that the brain’s resident immune environment, as well as peripheral immune cell infiltration, modifies neural circuits, synaptic transmission, and behaviour^23,25,26^. However, the neuroimmune correlates of adolescent social isolation remain obscure. Here, we used a chronic adolescent stress model, Post-Weaning Social Isolation (PWSI)^27–29^. We report marked sex differences in behavior patterns following 7 weeks of prolonged social isolation after weaning. Male mice exhibited increased aggression, whereas females exhibited reduced social interactions.

Furthermore, to understand the effects of PWSI-induced behavioral changes on the central immune system, particularly key brain regions such as the PFC and hypothalamus, we measured cytokines and their downstream signaling genes. Among the proinflammatory cytokines crucial for brain-immune communication, we observed changes only in IL6 and in associated signaling genes, specifically in the hypothalamus. IL6, being a multifunctional cytokine with context dependent pro and anti-inflammatory roles as well as non-immune CNS functions, is intriguing to explore in sex-specific behavioral outcomes and mental illness induced by social stress.

## Materials and Methods

### Animals

All experimental procedures involving live animals were approved by the Institutional Animal Ethics Committee (IAEC) of the CSIR–Institute of Genomics and Integrative Biology (Approval No.: IGIB/IAEC/16/Dec/2024/03). The IAEC is registered under the Committee for the Purpose of Control and Supervision of Experiments on Animals (CPCSEA), Department of Animal Husbandry and Dairying, Ministry of Fisheries, Animal Husbandry and Dairying, Government of India (Registration No.: 9/1999/CPCSEA,09/03/1999). Male and female Balb/c mouse offspring bred in the institutional animal facility were used for the study. Animals were group-housed (3–4 per cage) under specific-pathogen-free (SPF) conditions in individually ventilated cages (IVCs) maintained at 24 ± 2 °C on a 12-h light/dark cycle, with ad libitum access to food and water. All handling and procedures were performed in accordance with institutional guidelines.

### Post-Weaning Social Isolation procedure

Post-weaning social isolation (PWSI) was implemented as an established paradigm to elicit stress-induced behavioral phenotypes relevant to psychiatric disorder models. Weaning was conducted on postnatal day (P21), after which both male and female mice assigned to the PWSI cohort were housed singly in standard cages for five weeks; bedding and cages were replaced weekly to maintain hygiene without providing additional enrichment. Following this period, PWSI mice remained in single housing for an additional two weeks to sustain the isolation condition. After seven weeks of PWSI, animals were evaluated for offensive aggression using the resident–intruder (RI) assay. Age-matched control mice were maintained under standard group-housing conditions (3–4 per cage).

### Resident Intruder test

RI test for aggression was performed in male and female ‘Group Housed control’ mice who were not exposed to PWSI stress and ‘PWSI adult’ mice who were PWSI stressed based on the protocol reported earlier^30^. PWSI animals were kept in isolation for 7 weeks, while control animals were transferred to single housing for 72 h before RI testing to equilibrate immediate housing conditions without inducing long-term isolation effects. Each of these mice, referred to as “resident,” was exposed to two categories of unfamiliar intruders for 10 min per day for 2 consecutive days. Each day, the resident was introduced to a different intruder in the following manner: day 1 - same sex and 10% less body weight; day 2 - anaesthetized of the same sex and similar body weight. Of note, an anaesthetized intruder was incorporated in our study paradigm to assess pathological/maladaptive forms of aggression.

Behavioral measures including clinch attacks, approach responses, social investigation, anogenital sniffing, rearing, lateral threats, upright postures, pinning/keep-down, chasing, non-social exploration, and inactivity were analyzed as proportions of the total observation period. Attacks on vulnerable body parts, such as the neck, abdomen, and snout, were also recorded. Social exploration comprised the summed duration of social investigation, both self-and allo-grooming, and ano-genital sniffing. Non-exploratory activity was defined as the time during which the resident mouse remained active but did not investigate the intruder. Rest was defined as periods during which the resident mouse remained stationary without engaging in exploratory behaviors. Animal phenotypes were assigned according to standard ethological criteria reported in earlier studies^31,32^, with further details provided in the Results and Figure 1. Behavior was manually scored by a blind observer who was unaware of the animal identities and group assignments.

**Figure 1:**
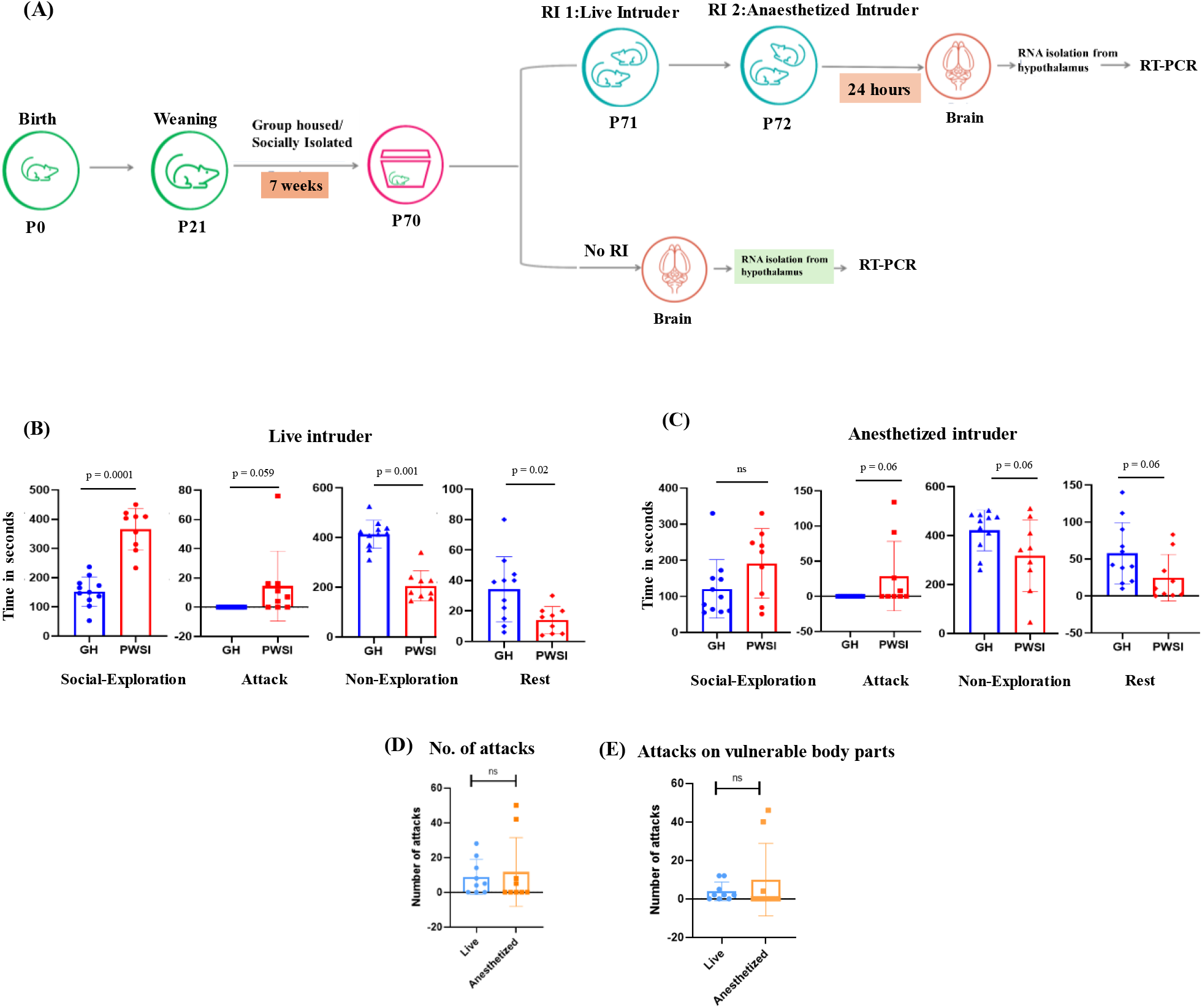
Escalated and maladaptive aggressive behavior by PWSI male mice. (A) Experimental Timeline of Post-Weaning Social Isolation (PWSI) procedure and Resident Intruder (RI) test. (B) Behavioral profiles of Grouped Housed (GH) and PWSI male mice towards live intruder and (C) anaesthetized intruder as measured by time spent in social exploration, attack, non-social exploration and rest or inactivity (D) Total number of attacks during the observation period and (E) number of attacks on vulnerable body parts by PWSI mice on both live and anaesthetized intruders. Data are presented as mean ± SD. ns-non-significant.

### RT-PCR

Briefly, total RNA was isolated from the hypothalamus and the prefrontal cortex (PFC) of mice using a conventional RNA isolation protocol. cDNA was synthesized from 2 µg of the total RNA using reverse transcription. RT-PCR was performed using SYBR Green Master Mix on a Bio-Rad CFX384 Real-Time System. All primers used for RT-PCR are listed in Supplementary Table 1. Quantification of each target was performed by normalizing it to the internal control, GAPDH.

**Table 1:**
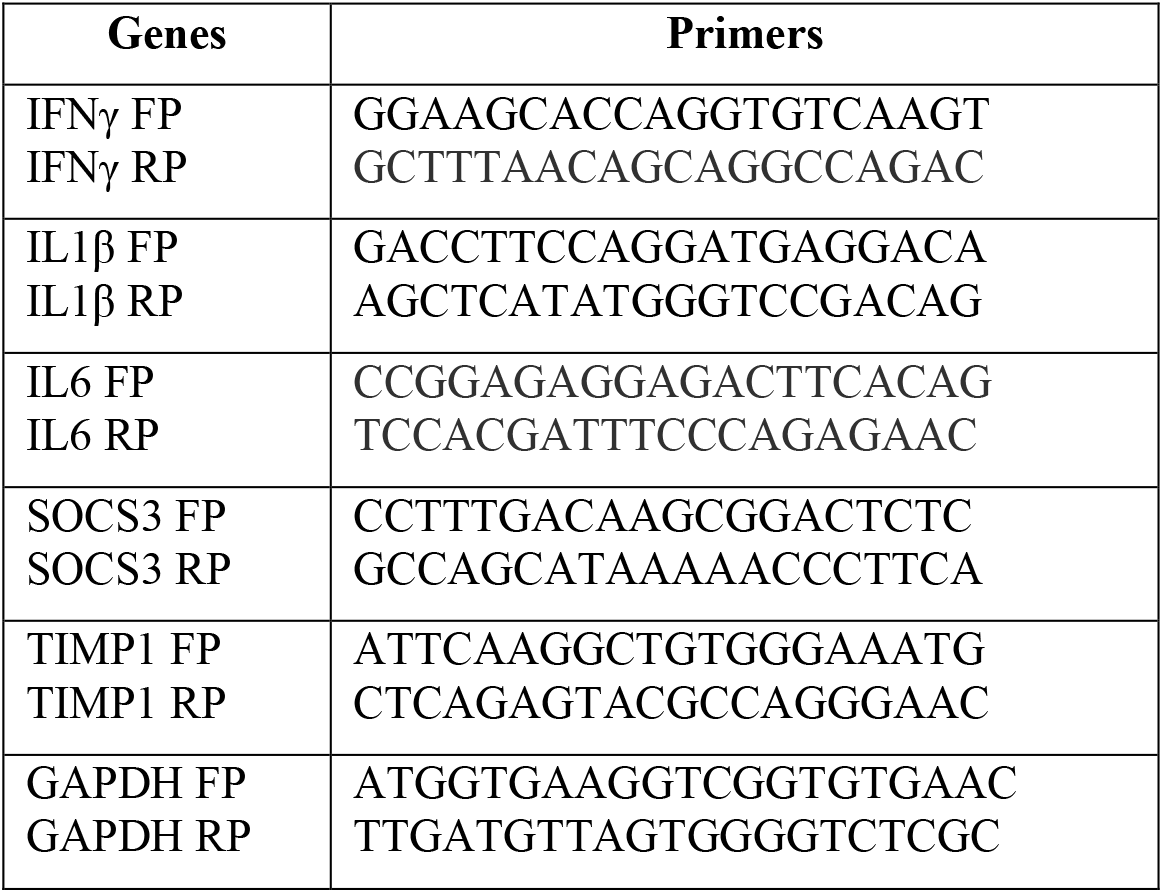
Primer sequences for RT-PCR.

## Results

### PWSI induced escalated and maladaptive aggressive behavior in male but not female mice

Behavioral encounter of Group-housed (GH) and PWSI resident males with live and anaesthetized intruders was assessed using the RI test of aggression (Fig. 1A). PWSI males spent more time in social exploration and showed markedly higher frequency of attacks on live intruders relative to GH controls (Fig. 1B). Mean time spent by GH mice was 152.6 and 0 seconds, while mean time spent by PWSI was 365.4 and 14.44 seconds in social exploration and attack, respectively. Differences between means (PWSI - GH) ± SEM for social exploration was 212.8 ± 27.21 seconds and for attack duration was 14.44 ± 7.18 seconds. PWSI males also attacked anaesthetized intruders (Fig 1C). Mean time spent in attack by GH males was 0 seconds, while mean time spent by PWSI males attacking the anaesthetized intruder was 28.78 seconds. The difference between mean time (PWSI - GH) ± SEM for attack duration was 28.78 ± 14.80 seconds. In addition to increased attack counts, PWSI mice targeted vulnerable body regions—including the neck, snout, and ventral areas—more frequently (Fig 1E). Aggression towards anaesthetized intruders was greater than that directed towards live intruders, indicating an escalation of offensive behavior (Fig 1D-E). Together, these findings demonstrate that PWSI induces a robust aggressive phenotype in male mice that is independent of intruder reactivity or behavioral cues, thereby generating “maladaptive or pathological aggression” in these mice.

Contrary to males, neither GH nor PWSI female mice showed any signs of offensive aggression towards intruders. However, PWSI female mice showed significantly reduced social exploration and increased inactivity/rest during the RI test sessions compared to GH controls. The mean time spent in rest by GH females was 39 seconds, while mean time spent by PWSI females in rest was 267.2 seconds. The difference between mean time (PWSI - GH) ± SEM for rest duration was 228.2 ± 21.49 seconds. Reduced social interaction coupled with inactivity might be indicative of social withdrawal, or a defensive stress coping strategy (Fig. 2B)

**Figure 2:**
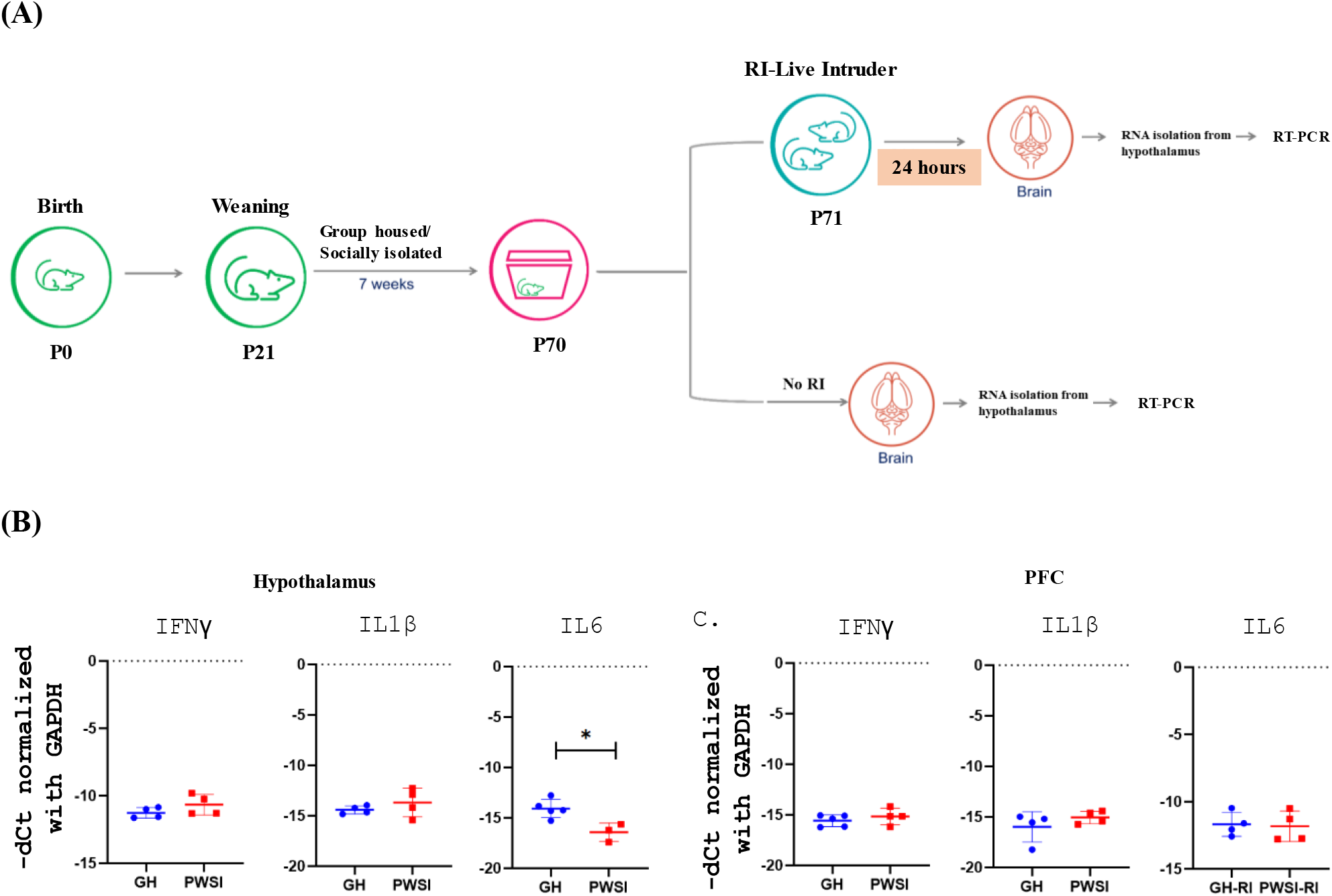
Non-aggressive and reduced social exploration behavior by PWSI female mice. (A) Experimental Timeline of Post-Weaning Social Isolation (PWSI) procedure and Resident Intruder (RI) test. (B) Behavioral profiles of Grouped Housed (GH) and PWSI female mice towards live intruder as measured by time spent in (i) social exploration, (ii) attack, (iii) non-social exploration and (iv)rest or inactivity. Data are presented as mean ± SD. ns-non-significant.

### IL6 and its responsive genes were downregulated in the hypothalamus of PWSI-induced aggressive male mice

We examined expression of cytokine mRNA levels in hypothalamus and PFC of male mice. Previous studies have established the role of hypothalamus in psychosocial stress response and reactive aggressive behaviour^33^. PFC acts as a top-down regulator of subcortical structures including hypothalamus and amygdala for expression of social behavior especially aggression and impulsivity^34^. We measured mRNA levels of pro-inflammatory cytokines IFN-γ, IL-1β, and IL6 in brain regions using RT–PCR. IL6 mRNA levels showed significant reduction in PWSI aggressive males as compared to GH controls (Fold Change: 0.006, 0.01, 0.23, 0.34; p < 0.05) whereas IFN-γ, IL-1β mRNA were unaltered (Fig. 3A). None of the cytokines showed significant differences in transcript levels between GH and PWSI males (Fig. 3B).

**Figure 3:**
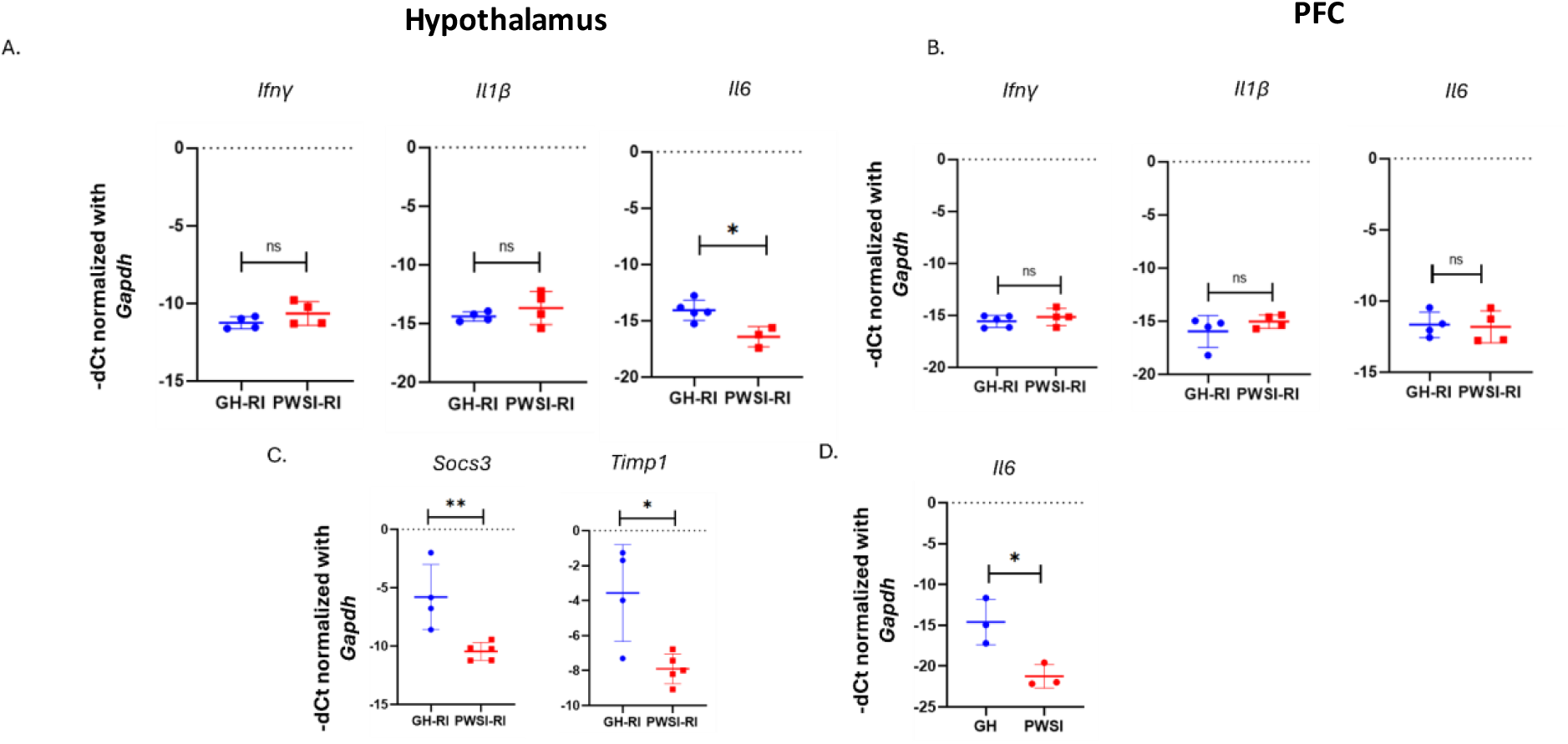
IL6 and its responsive genes (Socs3 and Timp1) were downregulated in the hypothalamus of PWSI-induced aggressive male mice. RT-PCR analysis of Ifnγ, Il1β, and Il6 mRNA levels in (A) hypothalamus, (B) PFC of GH and PWSI males after RI test (C) RT-PCR analysis of Socs3 and Timp1 mRNA levels in the hypothalamus of PGH and PWSI males after RI test (D) Il6 mRNA levels in GH and PWSI males without RI test. GH-Group housed; PWSI-Post-weaning Social Isolation; RI-Resident-Intruder test. Data are presented as mean ± SD* = p<0.05; ** = p<0.005, ns = non-significant.

Given that IL6 signaling regulates multiple downstream target genes, we subsequently examined the expression of the IL6-responsive genes Socs3 and Timp1 in the hypothalamus of male mice with PWSI-induced aggression. Both Socs3 and Timp1 transcript levels were significantly downregulated in PWSI-induced aggressive male mice (Socs3 FC: 0.002, 0.048, 0.002, 0.007; Timp1 FC: 0.0042, 0.0046, 0.107, 0.002, 0.068; p < 0.05) as compared to GH controls (Fig. 3C). Baseline IL6 mRNA levels in the hypothalamus of PWSI males without RI test were also reduced as compared to respective GH control males (Fig 3D). Whether this reduction in IL6 is mechanistically required for the emergence of aggressive behavior in PWSI males remains to be established.

### Expression of microglia and astrocyte markers downregulated in PWSI males in brain region-specific manner

The brain’s resident immune cells, astrocytes and microglia, are essential mediators of synaptic plasticity^35^, blood-brain barrier integrity^36,37^, and neuroinflammatory processes^38^. Previous studies have established that changes in the density and activation states of these glial cells directly influence behavioral outcomes^39^. More importantly Il6 is mainly secreted by astrocytes and microglia in the hypothalamus in response to immune challenges. Hence, we quantified mRNA levels of Iba1 (microglia) and Gfap (astrocytes) in the hypothalamus and PFC of PWSI-induced aggressive males. We observed downregulation of Iba1 mRNA levels in the hypothalamus (Fig. 4A) and Gfap mRNA levels in the PFC (Fig. 4B) in these mice, indicating region-specific changes in the glial marker expression.

**Figure 4:**
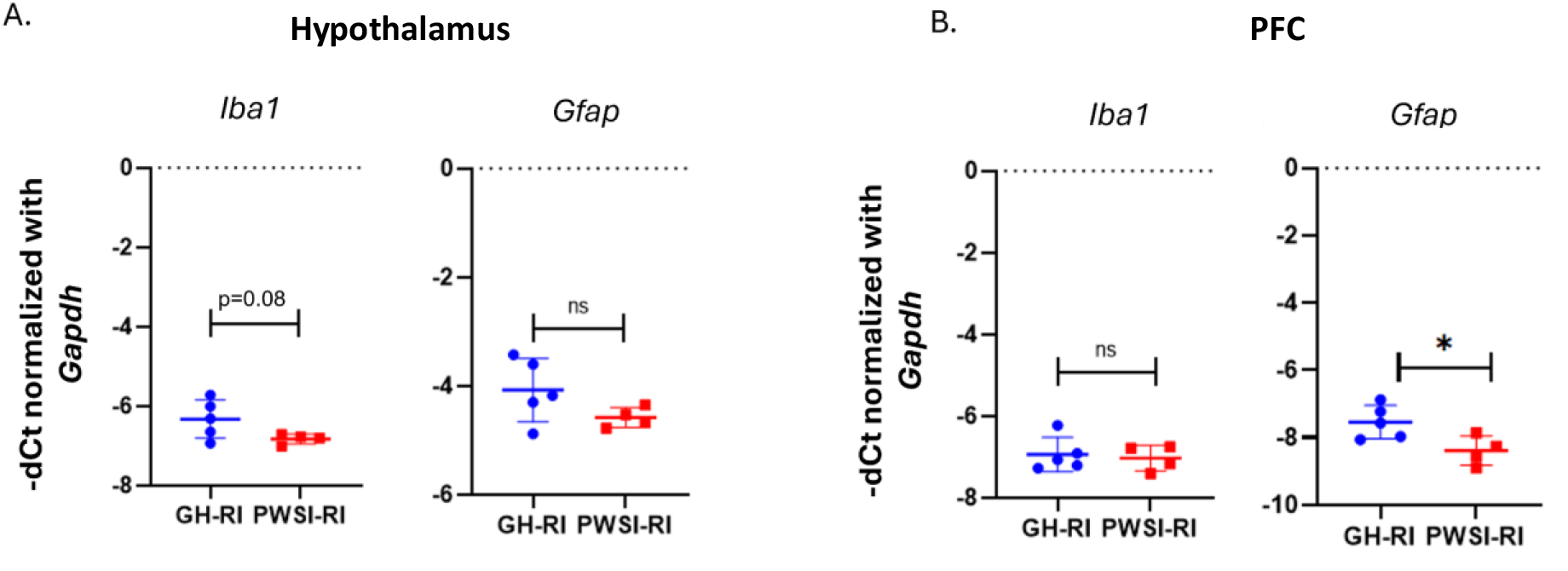
Brain region-specific downregulation of glial markers in PWSI induced aggressive males. RT-PCR analysis of Iba1 and Gfap mRNA levels in (A) hypothalamus, (B) PFC of GH and PWSI males after RI test GH-Group housed; PWSI-Post-weaning Social Isolation; RI-Resident-Intruder test. Data are presented as mean ± SD * = p<0.05, ns - non-significant.

### Hypothalamic IL6 downregulated in PWSI exposed females with quantitative sex differences

We performed RT-PCR analysis of cytokine mRNA (IFN-γ, IL-1β, and IL6) in the hypothalamus of PWSI-exposed females after the RI test. Despite sexual dimorphism in behavioral response, PWSI-exposed non-aggressive females also showed reduction in hypothalamic IL6 mRNA (FC: 0.441, 0.521,0.926,0.730) as compared to GH controls (Fig. 5A), similar to males. We further assessed the expression of the IL6-responsive genes and observed a trend of downregulation of Timp1 (FC: 0.107, 0.075, 0.342, 0.174) in the hypothalamus of female mice exposed to PWSI (Fig. 5B). Hypothalamic IL6 mRNA was also reduced in PWSI-exposed females not subjected to the RI test (Fig. 5C).

**Figure 5:**
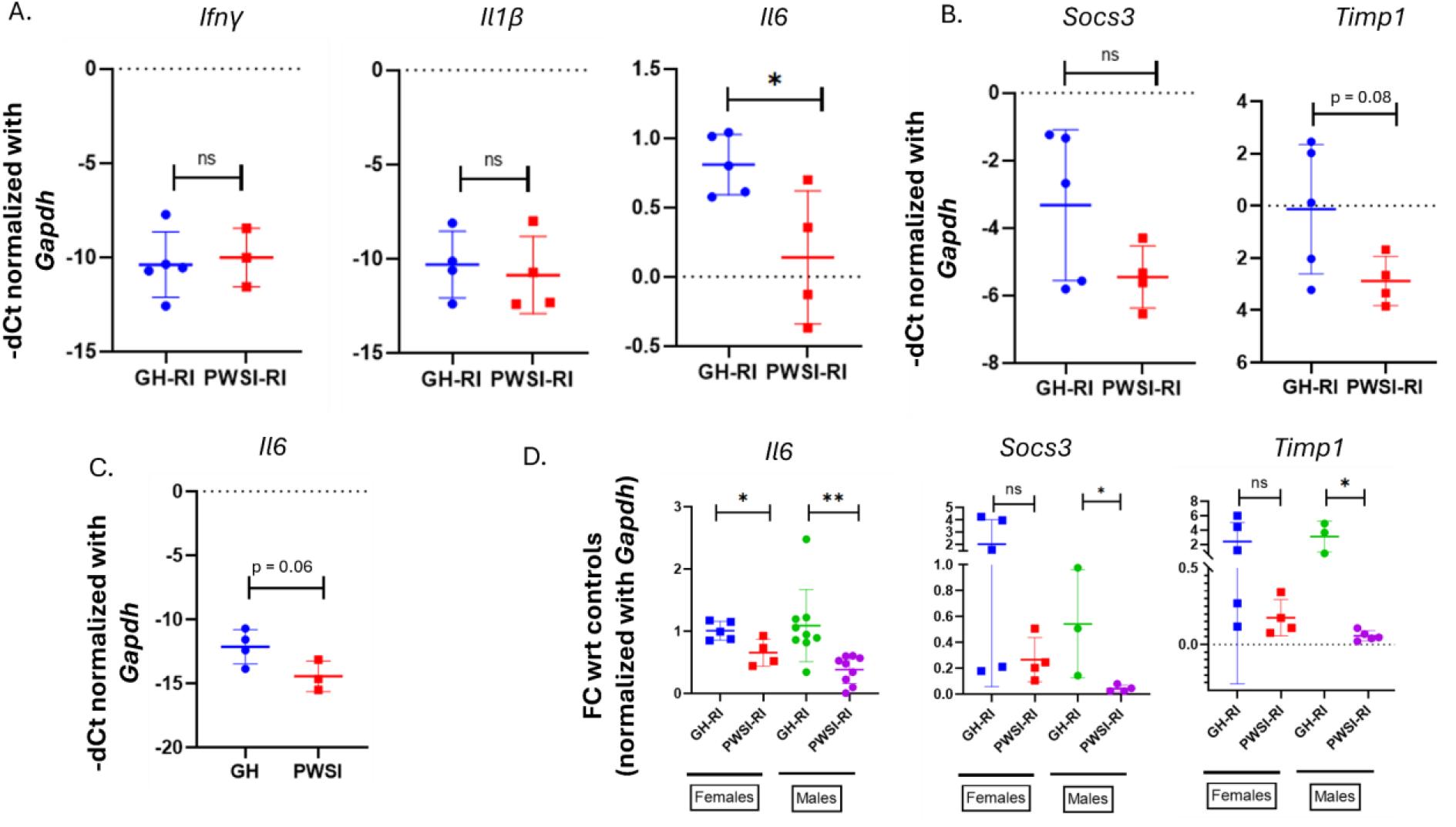
IL6 downregulated in the hypothalamus PWSI induced PWSI-exposed non-aggressive females. A. RT-PCR analysis of cytokines in the hypothalamus of PWSI-exposed female mice. B. RT-PCR analysis of Socs3 and Timp1 in the hypothalamus of PWSI-exposed female mice. C. RT-PCR analysis of *Il6* in PWSI male mice (without RI). D. Comparison of fold changes of Il6, Socs3, and Timp1 in PWSI-induced aggressive male, and PWSI-exposed non-aggressive female mice. GH = Group housed; PWSI = Post-weaning Social Isolation; RI = Resident-Intruder test. Data are presented as mean ± SD. * = p<0.05; ** = p<0.005, ns = non-significant

Sex differences can be qualitative, referred to as sexual dimorphism, or quantitative in terms of magnitude^40,41^. PWSI induced downregulation in hypothalamic IL6 and downstream targets (Socs3 and Timp1) in both sexes despite differential behavioral outcomes prompted us to assess if there are any quantitative differences. Notably, PWSI-induced aggressive males exhibited a more pronounced reduction in IL6, and IL6 responsive gene expression than PWSI-exposed non-aggressive females (Fig. 5D), suggesting a sex-dependent modulation of the hypothalamic IL6 signaling axis.

## Discussion

In this study, we demonstrated the potential role of hypothalamic IL6 in adolescent social isolation-induced aberrant behavior, particularly inter-male escalated aggression and its sex specificity. Previous studies have shown that social isolation leads to the development of aggressive behavior in rats^29^. Complementing these reports, our study shows that post-weaning social isolation induces aggression in male mice and furthermore reveals an a pathological phenotype characterized by short attack latency and biting of vulnerable body parts (neck, belly, and snout) in both live and anaesthetized intruders. Unlike males, female mice subjected to prolonged isolation from weaning exhibited no aggression toward same-sex intruders in adulthood. Instead, they displayed minimal social interaction, marked by increased resting or inactivity, indicative of either a reactive stress-coping strategy or low social motivation. Reduced interaction with intruders might also lead to territorial threats by resident females, though all these possibilities require testing using orthologous behavioral paradigms. Consistent with our findings, Tan et al. (2021) reported social withdrawal in female mice subjected to chronic social isolation stress^42^. However, in a previous study on rats by Oliviera et al., notable differences were observed in the development of aggression between the sexes^43^. The study showed the influence of neuropeptide systems involved in aggression regulation, notably affecting both the oxytocin and arginine-vasopressin pathways within major brain regions, including the hypothalamus^43^. Sex differences in social isolation-induced aggression might help us to understand how teenage boys and girls cope or succumb differently to bullying-induced isolation, parental neglect, and digital isolation due to excessive screen time, all of which need biological interventions.

Among pro-inflammatory cytokines, only Interleukin-6 (IL6) showed hypothalamus-specific downregulation in both sexes following PWSI and, more importantly, both prior to and 24h after a behavioral encounter with intruders. IL6 is a multifaceted cytokine secreted by immune cells during both innate and adaptive immune responses. It shows pro-inflammatory, anti-inflammatory, and non-immune functions in a context-dependent manner. Versatility of IL6 is mediated by multiple signaling cascades, primarily the JAK/STAT and NF-κB pathways^44,45^.

Beyond its role in neuroinflammation, IL6 plays important roles in CNS physiological processes, including neurogenesis, neuroprotection, and neurotoxicity^46–48^. In the context of Traumatic Brain Injury (TBI), IL6 has been observed to modulate microglial phenotype, thereby facilitating nerve repair and regeneration^49,50^. Moreover, during TBI repair, IL6 enhanced the survival of newly formed neurons. In neuropsychiatric systemic lupus erythematosus (NPSLE), a pathological condition in which neurons of the CNS and/or PNS are affected, IL6 induces apoptosis in adult hippocampal neural stem cells^51^. IL6 plays a dual role as both a protector of neurogenesis and an inducer of wound healing in diseases such as TBI^52^ and Alzheimer’s, while also contributing to blood-brain barrier (BBB) damage, thereby contributing to the pathophysiology of cerebral edema^52^. The dual nature of IL6 adds complexity to neuroimmune diseases, depending on factors such as the site of action, the duration and magnitude of its response, and the timing of its production^45^.

The hypothalamus is a major brain region involved in the regulation of stress response, aggressive behavior, motivation, energy homeostasis, and dietary regulation^53–58^. Under basal conditions, IL6 expression in the hypothalamus is relatively low and is primarily restricted to microglia, whereas IL6 receptors are widely expressed across multiple cell types, including neurons, astrocytes, microglia, and endothelial cells^59,60^. Beyond its established neuroinflammatory actions, hypothalamic IL6 contributes to a variety of physiological processes. It helps maintain stress response by directly activating the HPA axis, leading to the release of corticosterone^60^, and regulating the metabolism of tryptophan and serotonin^61^. IL6 has been implicated in the regulation of reward-seeking behavior and food intake within the lateral hypothalamus (LH), a key hypothalamic region that regulates hunger, thirst, arousal, and motivated behaviours^62^. Additionally, IL6 is produced by magnoneuronal neurons in the paraventricular (PVN) and supraoptic nuclei (SON) of the hypothalamus-neurohypophyseal system (HNS), identifying these neurons as an important non-immune source of circulating IL6 that regulates stress, metabolism and behaviour^60^. As a direct modulator of the HPA-axis^59^, IL6 plays a significant role in stress-induced anxiety^63^, psychosis^64^, depression^65^ and aggression^66,67^. Consistent with this, altered IL6 levels have been associated with several neuropsychiatric conditions, such as schizophrenia^68^ and substance abuse^69^. In the present study, Post-Weaning Social Isolation (PWSI) significantly down-regulated *Il6* expression in the hypothalamus of both male and female mice. This reduction in *Il6* transcript levels likely reflects the impact of prolonged isolation immediately after weaning, suggesting a functional role for hypothalamic IL6 in mediating responses to chronic early-life stress.

Activation of IL6 signaling induces the expression of several downstream target molecules, including Suppressor of Cytokine Signaling 3 (SOCS3) and Tissue Inhibitor of Metalloproteases 1 (TIMP1). Beyond its canonical role as a negative feedback regulator of cytokine signaling and inflammation^70^, SOCS3 has been implicated in chronic stress-induced activation of the HPA axis^71^ and in the pathophysiology of schizophrenia^68^. Notably, Meng et al. demonstrated that stress-induced SOCS3 expression within the paraventricular nucleus confers protection against inflammation-evoked pain^72^. TIMP1, another IL6-responsive molecule, is critical for maintaining the blood-brain barrier integrity^73^. Genetic deletion of TIMP1 in mice has been reported to produce anxiety-like behaviour^74^, underlining its potential relevance to stress-related neuropsychiatric phenotypes. Despite these observations, the relationship between PWSI, alterations in SOCS3 and TIMP1 levels, and behavioral outcomes has not been systematically investigated. In the present study, we provide the first evidence suggesting a potential involvement of SOCS3 and TIMP1 in the PWSI paradigm. To assess whether IL6 signaling is perturbed in PWSI-exposed mice, we quantified transcript levels of IL6-responsive genes, *Socs3* and *Timp1*. We observed a concomitant decrease of both transcripts in the hypothalamus. Taken together with the observed reduction in Il6 expression, these findings suggest that attenuation of hypothalamic IL6 signaling may contribute to the behavioral alterations induced by early-life social isolation. However, the specific neural circuits and molecular mechanisms linking PWSI-induced IL6 signaling deficits to the behavioral phenotype remain to be elucidated and warrant investigation.

Previous studies have identified microglia as the primary source of IL6 in a neuroinflammatory context^59^. Moreover, alterations in microglial abundance or activation state are known to modulate IL6 levels^75^, which can subsequently change the behaviour^76,77^. This suggests that PWSI may be driving changes in microglial levels or activation status, thereby reducing *Il6* levels. The potential interplay between decreased hypothalamic *Iba1* and decreased *Gfap* levels in the PFC, and how these changes contribute to diminished hypothalamic *Il6* signaling and behavioral outcomes in PWSI male mice, warrants investigation.

Interestingly, our findings demonstrate that PWSI produces sexually dimorphic behavioral outcomes in adulthood. Male mice exposed to PWSI exhibited pronounced pathological aggressive behaviour, whereas female mice were resilient to PWSI-induced aggressive behavior. Nonetheless, female mice displayed an alternative behavioral alteration, which could be observed as an increase in the time spent in the resting state during the RI test, indicative of impaired social interaction or reduced social engagement. Despite these sex-specific behavioral phenotypes, we observed a shared molecular signature in the hypothalamus of both males and females, characterized by reduced expression of *Il6* and its downstream responsive genes. This prompted us to investigate whether the magnitude of the molecular change differed between sexes. Notably, the magnitude of reductions in *Il6, Socs3*, and *Timp1* levels was substantially greater in aggressive male mice than in non-aggressive female mice. These data suggest that while the IL6 pathway disruption is a common consequence of PWSI, the extent of signaling attenuation may contribute to divergent behavioral outcomes between sexes. Prior work has highlighted sex-dependent functions of Lateral Hypothalamic (LH) IL6 in regulating motivated behaviors, with IL6 reported to increase food intake in males while enhancing food-motivated reward behavior in females^62^. In this context, the behavioral divergence observed in our model may reflect differences in neural circuits, neurochemical identity, or neuroanatomical targets among IL6-expressing neurons engaged by early-life social isolation. It is also plausible that sex-specific hormonal or developmental factors modulate the functional consequences of reduced IL6 signaling.

Overall, our results support a model in which hypothalamic IL6 signaling is broadly sensitive to early life social stress in both sexes, but sex-dependent network-level mechanisms shape the resulting behavioral phenotypes. Future studies are required to delineate the precise neuronal population and circuit dynamics through which reduced IL6 signaling differentially influences male and female behavioral outcomes following PWSI.

